# Analysis of Curcumin and Piperine for Breast Cancer Prevention and Treatment via Simulation and Artificial Intelligence Methods

**DOI:** 10.1101/2024.11.15.623750

**Authors:** Ashwin Sivakumar, Rishi Senthil Kumar, Sheena Christabel Pravin, Reena Monica P

**Affiliations:** School of Electronics Engineering, Vellore Institute of Technology, Chennai,India; MedxAI Innovations, Vellore Institute of Technology, Chennai,India

**Keywords:** Curcumin, Piperine, Neural Network, Molecular Docking, Dose-Response Curve

## Abstract

From published literature, the combined properties of Curcumin and Piperine infers that the two molecules are potential preventative agents. The nutraceutical combination of Curcumin and Piperine may aid in breast cancer prevention by enhancing Curcumin’s anti-inflammatory and antioxidant effects. In this paper, we demonstrate the following analysis on these two nutraceutical drug molecules in the effort to replicate these inferences in-silico: Drug Synergy Analysis, Molecular Docking and Simulation of Dose-Response curves. We have predicted the synergy (i.e., the effect of two drug molecules when administered simultaneously) between the two molecules on the basis of the Bliss’ Additivity Score. These scores have been predicted by devising a deep learning model using neural networks to determine that the two molecules of interest are synergetic in nature (i.e., one molecule improves the effectiveness of the other). The data used for this model has been extracted from the DrugComb database and filtered to having drug combinations under the MCF7 cell line. The drug molecules are converted from its original SMILES representation to the ECFP6 molecular fingerprint format prior to training the data with the output feature as the Bliss’ score. In this work we also study the interactions between the molecules of interest and protein targets which are of interest (TRPV1 and STAT3) via Molecular Docking. From this, we have been able to determine the binding affinity of the two ligands of interest by docking simultaneously. We have also identified the coordinates of the binding sites of the respective ligands and other potential binding sites for effective binding. From the binding free energies simulated from molecular docking, we have derived the IC50 values using the Cheng-Prusoff Equation. From these IC50 values we have derived the monotherapy dose-response curves with respect to the activation of each the protein targets and the molecules of interest over a range of molar concentrations (0.01um to 100um) using the 4 – parameter logistic curve equation. Furthermore, we have devised a 3D heatmap indicating the combined effectiveness of Curcumin and Piperine from the 4 – parameter log – logistic curve of the Loewe’s Additivity Model over each of the protein targets in focus.

## 1 Introduction

Curcumin and piperine, two natural compounds with notable bioactive properties, have emerged as potential adjuncts in the prevention and treatment of breast cancer. Curcumin, a polyphenolic compound derived from the spice turmeric (Curcuma longa), has been extensively studied for its anticancer properties [1]. Piperine, an alkaloid from black pepper (Piper nigrum), is often used in conjunction with curcumin due to its ability to enhance curcumin’s bioavailability, which otherwise suffers from poor absorption in the human body [2]. Together, curcumin and piperine offer promising therapeutic effects that could support conventional breast cancer therapies [3].

The lengthy and costly nature of drug discovery and analysis in the pharmaceutical industry, particularly during experimental phases, underscores the need for more efficient methods. Cheminformatics, combined with AI-driven techniques, is essential for efficiently screening drug targets and simulating drug candidates, significantly reducing both time and expenses.This proves to be a more efficient method to screen and validate claims in-silico [4]

This paper demonstrates a pipelined methodology that integrates machine learning and deep learning models to screen potential drug candidates and predict critical metrics such as drug synergy, response, and sensitivity. By combining predictive modeling with additional stages for molecular docking analysis, dose-response curve predictions, and ADME (Absorption, Distribution, Metabolism, and Excretion) predictions, this pipeline enhances the drug discovery process with comprehensive, data-driven insights. Ultimately, this approach streamlines drug development, enabling more accurate identification of effective drug combinations and supporting the creation of targeted, efficient treatments for complex diseases.

## 2 Literature Review

### 2.1 Experimental Review of Curcumin and Piperine

Curcumin exhibits a broad spectrum of anticancer activities through various molecular mechanisms, primarily involving inhibition of inflammation, induction of apoptosis, and suppression of tumor cell proliferation [5]. One of the key pathways it influences is the nuclear factor-kappa B (NF-kB) signaling pathway, a pathway known to promote cancer cell survival and proliferation. By inhibiting NF-kB, curcumin can potentially reduce tumor growth and decrease cancer cell resistance to chemotherapy [6]. Additionally, curcumin downregulates the expression of multiple pro-inflammatory cytokines, which are often elevated in breast cancer patients and contribute to tumor progression [1].

Curcumin also exerts influence on cellular apoptotic processes by modulating proteins such as p53 and Bax, which are critical regulators of programmed cell death. This effect is particularly significant in hormone-responsive breast cancer, where tumor growth depends on signals from estrogen receptors. Curcumin’s ability to downregulate estrogen receptors may help reduce the proliferation of hormone-sensitive breast cancer cells [3]. Furthermore, curcumin has been shown to inhibit angiogenesis—the formation of new blood vessels that supplies nutrients to tumors, thereby starving cancer cells and slowing tumor growth [7].

Piperine is known to improve curcumin’s bioavailability significantly, with studies showing that it can increase absorption by as much as 2000% [6]. The enhanced bioavailability is attributed to piperine’s ability to inhibit enzymes responsible for drug metabolism, thus preventing the rapid breakdown of curcumin in the liver and intestines [8]. This synergy makes curcumin-piperine combinations more effective in therapeutic settings compared to curcumin alone.

In breast cancer specifically, piperine not only enhances curcumin absorption but also exhibits its own anticancer properties. It has been reported to inhibit breast cancer cell proliferation by inducing cell cycle arrest and apoptosis. Piperine’s mechanisms include the modulation of several molecular targets, including enzymes and proteins that regulate cancer cell growth [2]. Additionally, piperine may increase the sensitivity of breast cancer cells to chemotherapy, potentially reducing the required dose of chemotherapeutic agents and minimizing side effects [5].

The combination of curcumin and piperine has shown promise in preclinical studies, suggesting potential for integrative cancer treatment [7]. Studies indicate that this combination may work synergistically with conventional therapies, potentially enhancing the effectiveness of chemotherapy and radiation while reducing their adverse effects [8]. This could be particularly valuable in cases where breast cancer patients experience resistance to standard treatments or have difficulty tolerating high doses of chemotherapeutic drugs [6].

### 2.2 Simulation and Use of Artificial Intelligence for in-silico analysis

The use of graph neural networks has been proven useful for generating new drug combinations. Neural networks such as MLP Regressor is further used to find drug synergy classifications [9]. Another variation of graph neural networks called DDoS has been implemented with the objective of predicting drug synergy [10]

The synergy of drug combinations can be categorized into 3 types: Synergistic, Antagonistic and Additive. This is based on types of scoring methods (HSA, Bliss, Loewe’s etc). For our simulations we consider the Loewe’s Additivity Model. [11] When drugs are administered simultaneously to a patient, their effects can be either amplified or diminished, potentially leading to side effects. These types of interactions are known as drug-drug interactions (DDIs). Predicting potential DDI helps reduce unanticipated drug interactions and drug development costs and optimizes the drug design process.[12, 13] Representation of molecules can be done in various approaches, one of the descriptors is done via molecular fingerprinting techniques such as ECFP and Morgan Fingerprinting [14]. Image representation techniques can also be applied such as ImageMol [15].

## 3 Methodology

### 3.1 Prediction of Drug Synergy using Neural Networks

The dataset required for the prediction is derived from the DrugComb database [16]. The query given to the database API filters out the necessary drug combinations from the MCF7 cell line. Combination only existing of a single drug were removed during data preprocessing. The extracted dataset consists of the SMILES representation of the drug combinations [17], the corresponding concentrations and the respective drug synergy scores (ZIP, BLISS, Loewe’s and HSA) and each drug’s IC50 values.

Using RDKit [14], we convert all the SMILES of the drug molecules into a quantifiable format. In this case we convert it into the ECFP6 Fingerprint [18]. This is a version of morgan fingerprinting. The SMILES is converted into a single 300 bit string consisting of 0s and 1s. The 1 in a particular bit position represent a particular substructure which extends to 6 atoms in length and 0 represents the absence of that particular substructure. For each drug combination ECFP6 fingerprint, each bit is split into a feature vector.

The target feature for the classification model is modified on the basis of the additive, synergistic and antagonistic threshold of each drug synergy score. For example, the Loewe’s score is considered additive if it lies between -5 to 5. Above 5 is considered synergistic and below -5 is considered antagonistic. Similarly, for bliss model we consider -0.1 to 0.1 as additive. Scores above 0.1 would be considered synergistic and below -0.1 is considered antagonistic.

The dataset is trained into a feed - forward neural network. We have used 4 Dense hidden layers of 1024, 512, 256, 128 nodes each. The ReLU activation function is used as the primary activation function [19].

Both models have been created using Python and PyTorch [20]

### 3.2 Determining Binding Affinity and potential binding sites by Molecular Docking

The SMILES (Simplified Molecular Input Line Entry System) representation [17] of Curcumin [21] and Piperine [22] is attained from PubChem. By entering the SMILES representation of both the ligands in OPENBABEL [23] as input, the Protein Data Bank (PDB) format files are attained. From the protein data bank [24], the PDB files of TRPV1 and STAT3 are retrieved. The PDB IDs for TRPV1 and STAT3 are 3J5P and 6NUQ respectively.

We used ChimeraX [25, 26] to prepare the protein receptor for docking. First, we removed all Non-standard residues: Select—Residues—All Nonstandard, Action—Atoms/Bonds—Delete. Then, we prepared it for docking: Tools—Structure Editing—Dock Prep. We did this for both protein receptors. To perform molecular docking, we used PyRx Software [27]. First, we loaded both receptors’ PDB files and then converted them into a PDBQT file. After that, we loaded the two ligands as PDB files and converted them into PDBQT files too. We performed molecular docking of both ligands at the same time with each protein separately.

We used FPocket software [28] to get more information about the binding site of the two ligands. From FPocket we were able to get binding site data like the number of alpha spheres, Solvent Accessible Surface Area (SASA), Volume, etc.

To visualize the docking, we first downloaded the mode 0 docking orientation of both the ligands and the receptors as PDB files from PyRx. Then, in ChimeraX we loaded the receptor’s PDB file and the ligand’s mode 0 orientation PDB file to visualize the docking and to find the binding site.

### 3.3 Modeling of Dose Response Curves

In this section we calculate the dose response curve. We determine over a particular range of concentrations, the percentage inhibition activity of the of Curcumin and Piperine both independently as well as in a combined manner over the set of neuronal modulators as mentioned previously. In order to chart out these models, there are certain derivations and assumptions to be considered.

Firstly, we retrieve the binding affinity free energies from molecular docking and find the inhibition constant *K*_*i*_. The formula is given as Δ*G* is equal to N times natural log of *K*_*i*_ divided by C. Here, R stands for the universal gas constant. For reference, we have taken the room temperature as 298K and and C is taken as the reference concentration at 1M.

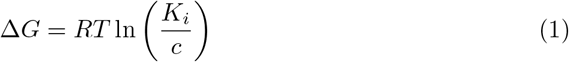

From the Inhibition constant, we the *IC*_5_0 value, that is the concentration at which the inhibition of our target hormone or protein is 50%. We determine this using the Cheng - Prusoff equation [29]. The main assumption we take from this equation is that we assume that the concentration of maximum substrate saturation is equal to the Michelis-Menten Constant. This shows that the inhibition constant *K*_*i*_ is directly related to the inhibitor’s effectiveness at half maximum substrate saturation. Here, *K*_*i*_ represents the affinity of the inhibitor of the enzyme, indicating that the inhibitor has a significant impact at this substrate concentration. This way we reduce the equation such that twice the inhibition constant is equal to the *IC*_5_0 value.

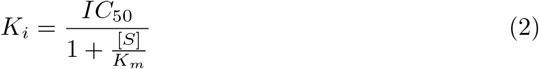

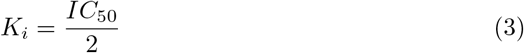

In order to model the dose response curve, we use a 4 parameter logistic curve. In this *Y*_*min*_ and *Y*_*max*_ stands for the percentage inhibition where *Y*_*max*_ is 100% and *Y*_*min*_ is 0% x stands for the range of concentrations. Here we assume it from 0.01 micromolar up to 100 micromolar [16].

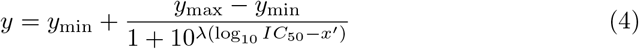

In order to find the combined effect of both the drugs administered simultaneously. We have devised a 3D heat map. Using the four parameters log logistic curve from the Loewe’s additivity model. Here m1 and m2 represent the IC50 values [16, 30, 31]

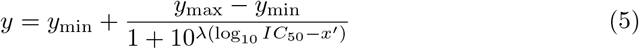

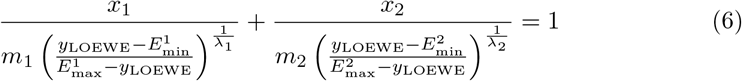

The plotting of monotherapy curves and the 3D heatmap have been carried out using MATLAB [32]

## 4 Results

### 4.1 Prediction of Drug Synergy using Neural Networks

For our regression model we keep the Bliss synergy score as our target feature. We have run the model at a benchmark of 300 and 500 epochs. The Bliss score is predicted to be 1.6826 for 300 epochs and 1.6555 for 500 epochs. As the two predicted score satisfy the synergy threshold (Score*>*0), hence, we can predict that the drug combination is synergetic in nature. We further validate this claim with our classification model. Table 1 shows the metrics of our proposed regression model: Mean Squared Error (MSE), Root Mean Squared Error (RMSE), Mean Absolute Error (MAE) and *R*^2^ value. Figure 4 shows the predicted versus the actual values plotted for our test dataset.

**Table 1.** Regression Model Performance Metrics at Epochs 300 and 1000.

|  | Epoch 300 | Epoch 1000 |
| --- | --- | --- |
| **MSE** | 0.3162 | 0.2914 |
| **RMSE** | 0.5623 | 0.5398 |
| **MAE** | 0.3386 | 0.2910 |
| **$R^2$ Value** | 0.9867 | 0.9878 |
| **Predicted Score** | 1.6826 | 1.6555 |

**Fig. 1.**
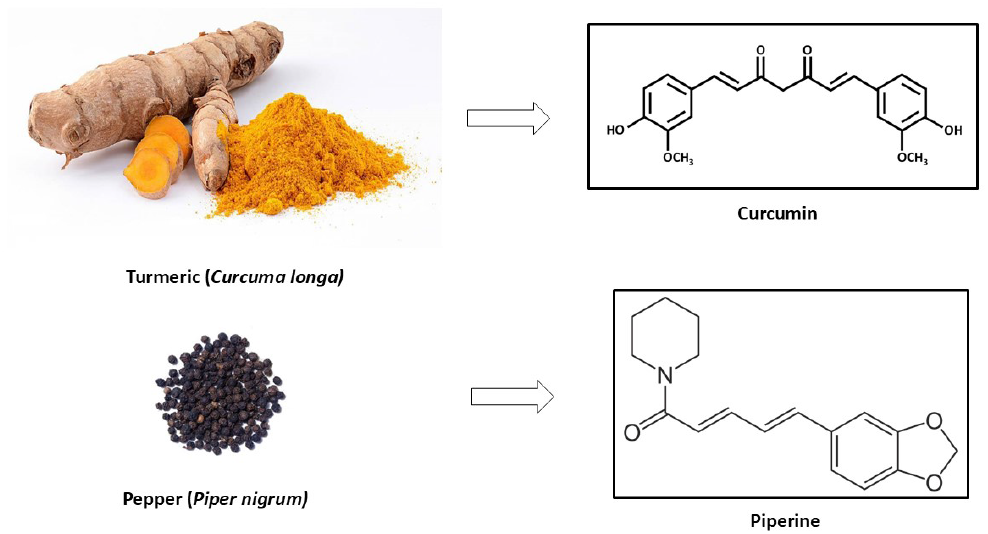
Curcumin derived from Turmeric(*Curcma longa*) and Piperine derived from Pepper(*Piper nigrum*)

**Fig. 2.**
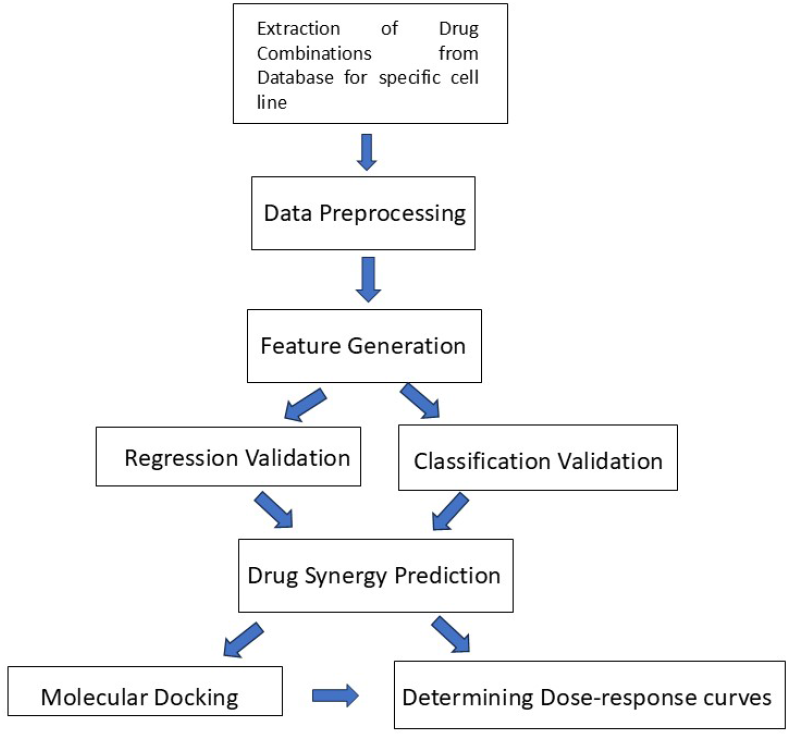
Graphical illustration of methodology for in-silico analysis of Curcumin and Piperine

**Fig. 3.**
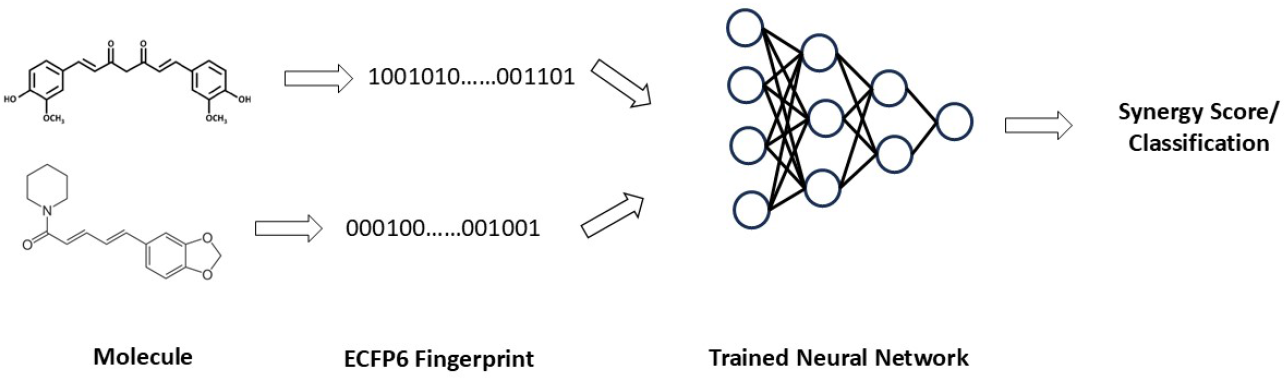
Graphical illustration of Prediction and Classification of Drug Synergy Scores

**Fig. 4.**
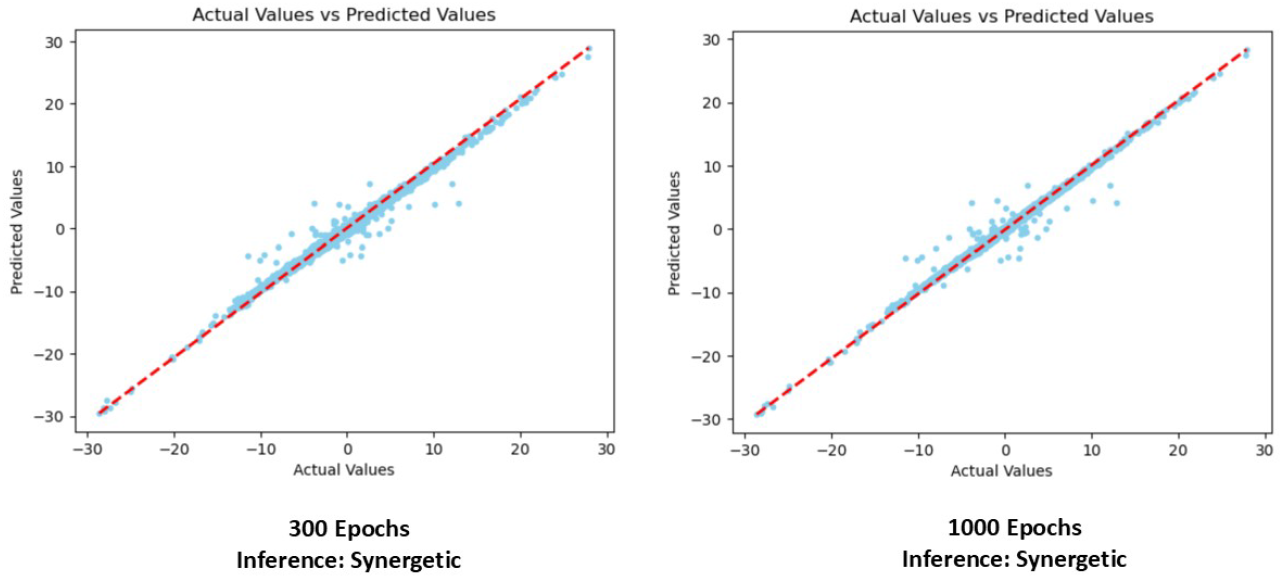
Predicted versus actual values plotted for 300 and 500 epochs show linearity. Inference of our required drug combination is also specified as synergetic in both cases.

For our classification model, we modify the Bliss synergy score into three categories. -1 for antagonistic, 0 for additive and 1 for synergistic. We have run the model at a benchmark of 300 and 500 epochs. In both cased the model predicted that the drug combination is synergistic in nature. Table 2 shows the metrics of our proposed classification model: Precision, Recall, F-1 score and Accuracy of the model. Figure 5 shows the confusion matrix representing the accuracy of our model for our test dataset.

**Table 2.** Classification Model Performance Metrics at Epochs 300 and 1000.

| Class | Precision | Recall | F1-score | Accuracy | Epochs |
| --- | --- | --- | --- | --- | --- |
| Class -1 | 1.00 | 0.99 | 1.00 | 0.9955 | 300 |
| Class 0 | 0.99 | 0.98 | 0.99 |  |  |
| Class 1 | 0.99 | 1.00 | 1.00 |  |  |
| Class -1 | 0.99 | 1.00 | 1.00 | 0.9955 | 1000 |
| Class 0 | 0.98 | 1.00 | 0.99 |  |  |
| Class 1 | 0.98 | 0.99 | 0.99 |  |  |

**Fig. 5.**
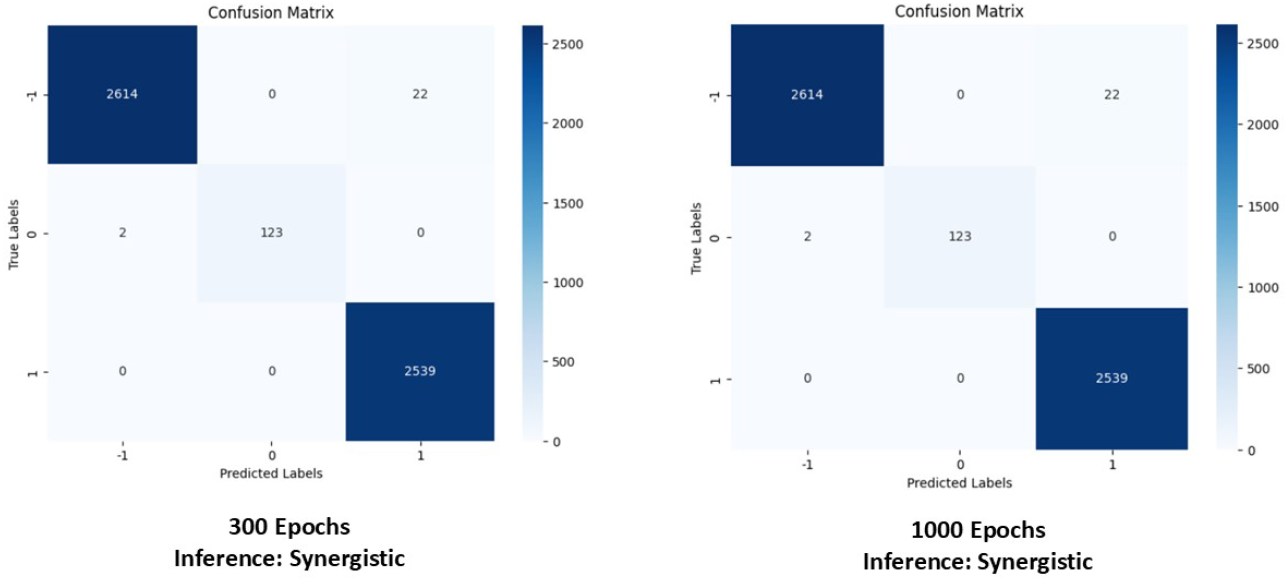
The confusion matrix represents the accuracy of the classification for 300 and 1000 epochs. Inference of our required drug combination is also specified as synergistic in both cases.

**Fig. 6.**
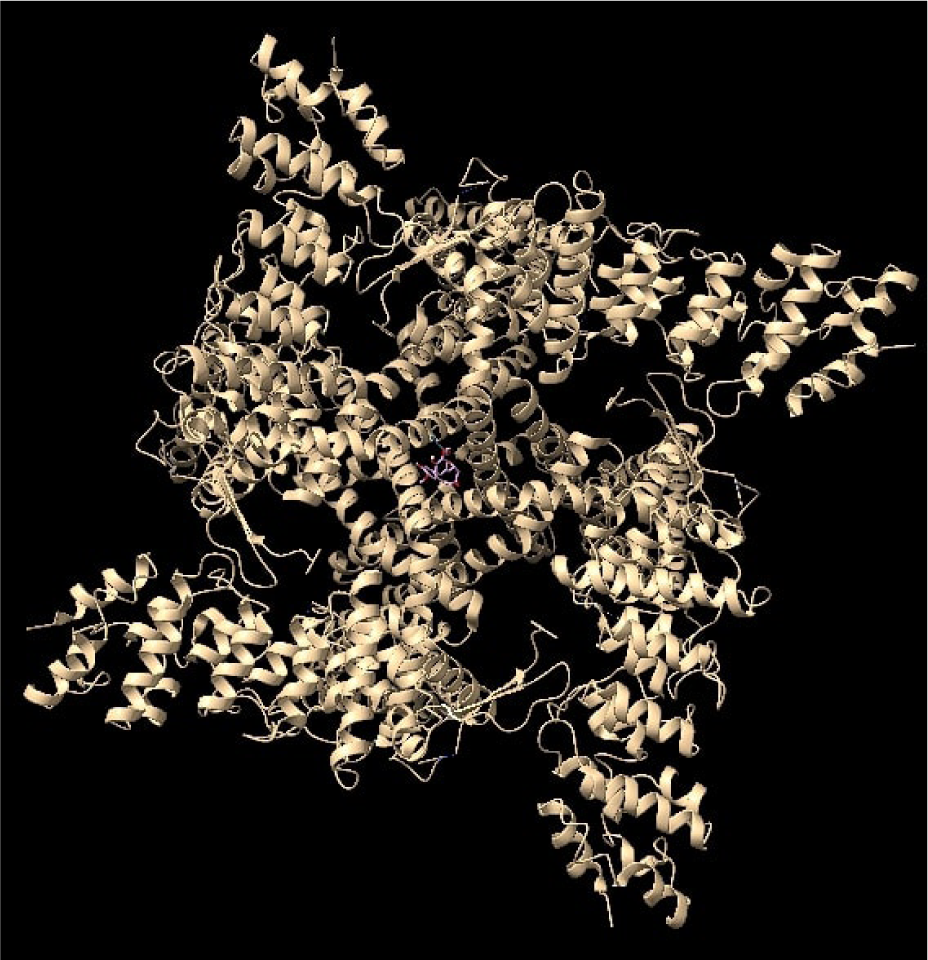
Simultaneous Docking of Curcumin and Piperine on TRPV1 Protein

**Fig. 7.**
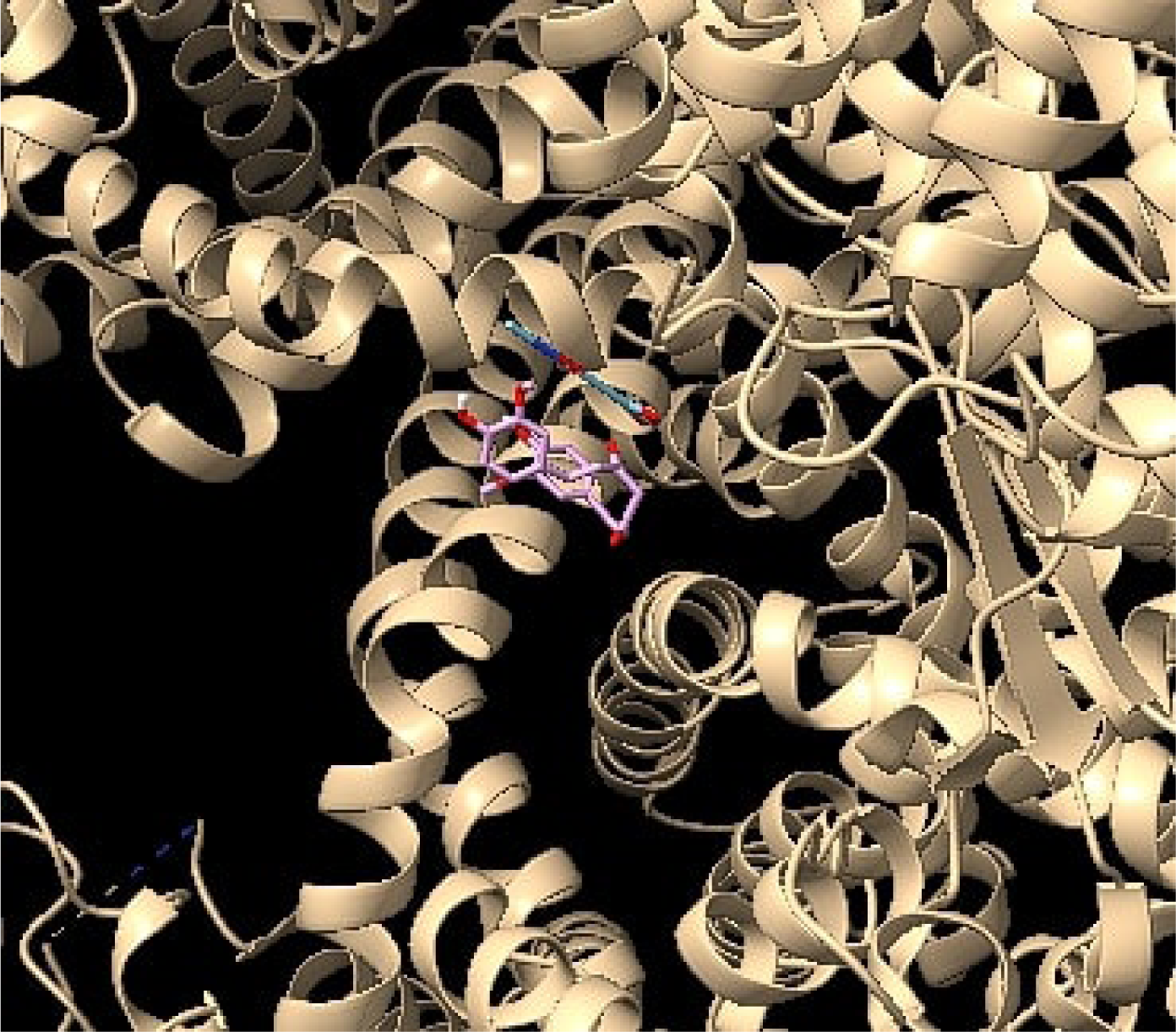
Close-up version of Figure 6 binding site. Lavender represents Piperine and Cyan represents Curcumin

### 4.2 Molecular Docking to determining Binding Affinity

Piperine had a strong binding affinity with both TRPV1 and STAT3. At the most optimal orientation (mode 0), Piperine had a binding affinity of -9.4 kcal/mol and - 10.6 kcal/mol with TRPV1 and STAT3 respectively. High binding affinity means that Piperine has a good potential of becoming a drug. High binding affinity also indicates that Piperine will target these proteins strongly and not attack normal functional proteins, which may cause drug side effects. At mode 0, Piperine had a root mean squared deviation of 0 in both the upper and lower bound.

Meanwhile, Curcumin showed a moderate binding affinity with both TRPV1 and STAT3. Even at the most optimal orientation, Curcumin only had a binding affinity of -6 kcal/mol and -6.2 kcal/mol with TRPV1 and STAT3. These binding affinity scores are low when compared to those of Piperine. At mode 0, Curcumin had a root mean squared deviation of 0 in both the upper and lower bound.

Table 3 and 4 shows the binding affinity of Piperine and Curcumin with respect to the TRPV1 and STAT3 protein respectively from mode 0 to 8.

**Table 3.** Binding affinity of Piperine and Curcumin on the inhibition of TRPV1.

| Ligand | Binding Affinity | rmsd/ub | rmsd/lb |
| --- | --- | --- | --- |
| Piperine Mode 0 | -9.4 | 0 | 0 |
| Piperine Mode 1 | -9.3 | 5.985 | 5.849 |
| Piperine Mode 2 | -9.2 | 8.327 | 7.372 |
| Piperine Mode 3 | -9.0 | 7.652 | 6.002 |
| Piperine Mode 4 | -8.8 | 6.036 | 4.03 |
| Piperine Mode 5 | -8.8 | 5.485 | 4.182 |
| Piperine Mode 6 | -8.8 | 6.217 | 4.832 |
| Piperine Mode 7 | -8.8 | 5.324 | 4.738 |
| Piperine Mode 8 | -8.4 | 7.806 | 6.006 |
| Curcumin Mode 0 | -6.0 | 0 | 0 |
| Curcumin Mode 1 | -5.3 | 2.321 | 1.502 |
| Curcumin Mode 2 | -5.3 | 3.081 | 1.972 |
| Curcumin Mode 3 | -5.1 | 10.741 | 5.646 |
| Curcumin Mode 4 | -5.1 | 11.392 | 7.003 |
| Curcumin Mode 5 | -5.0 | 8.624 | 6.486 |
| Curcumin Mode 6 | -5.0 | 3.113 | 2.072 |
| Curcumin Mode 7 | -4.9 | 9.357 | 5.674 |
| Curcumin Mode 8 | -4.8 | 9.848 | 6.363 |

**Table 4.**
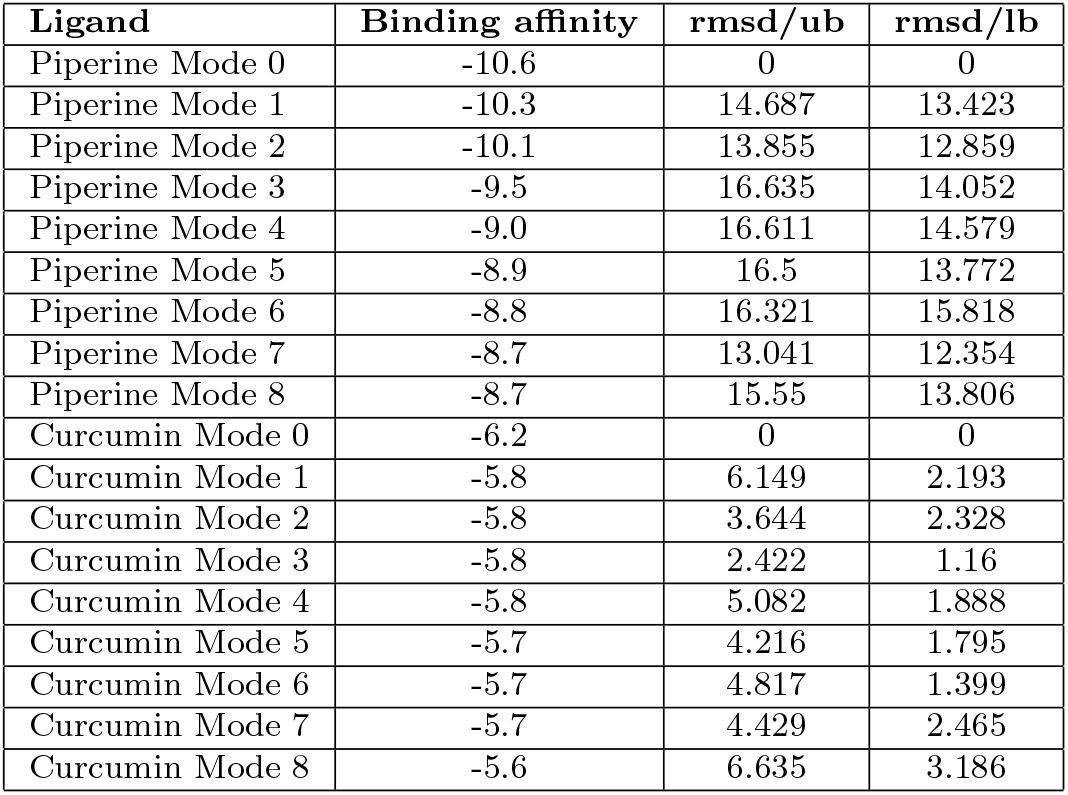
Binding affinity of Piperine and Curcumin on the inhibition of STAT3 protein.

From FPocket, we got to know that the binding sites in both protein receptors are quite spacious. The binding sites in TRPV1 and STAT3 have 112 and 60 alpha spheres respectively. With 112 alpha spheres, TRPV1 has a total volume of 1160.410 angstroms cube and a total SASA of 388.083 angstroms square. Meanwhile, STAT3 has a total volume of 559.860 and a total SASA of 151.023. When compared with TRPV1, STAT3 has relatively low volume and SASA but still is large enough to dock the two ligands.

### 4.3 Simulated Dose Response Curves

From molecular docking, we assume the binding affinity values of Curcumin and Piperine with respect to the STAT3 and TRPV1 proteins in the most stable state (Mode 0). This affinity value is considered as the binding free energy in equation 1. With respect to the TRPV1 protein, the inhibition constant values calculated from equation 1 are 0.1276 µM and 39.76 µM for Piperine and Curcumin respectively. Their respective IC50 values are calculated to be 0.2552 µM and 79.52 µM.

With respect to the STAT3 protein, the inhibition constant values calculated from equation 1 are 0.0168 µM and 28.368 µM for Piperine and Curcumin respectively. Their respective IC50 values are calculated to be 0.0336 µM and 56.736 µM from equation 2 and 3.

The Monotherapy Curves of Curcumin and Piperine with respect to the inhibition of TRPV1 and STAT 3 protein are illustrated in Figures 10 and 11. These plots are derived from the 4-parameter logistic curve from equation 4.

**Fig. 8.**
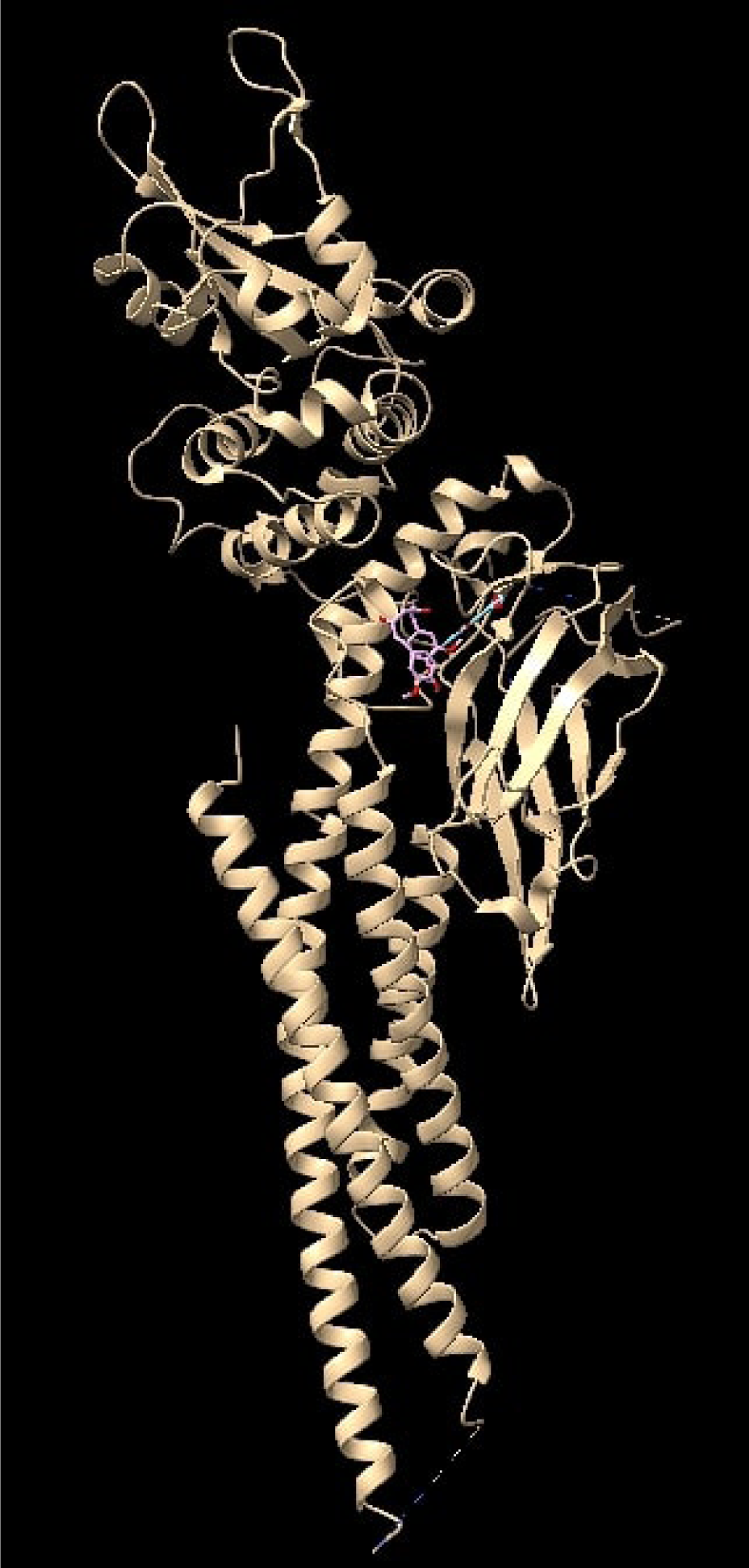
Simultaneous Docking of Curcumin and Piperine on STAT3 Protein

**Fig. 9.**
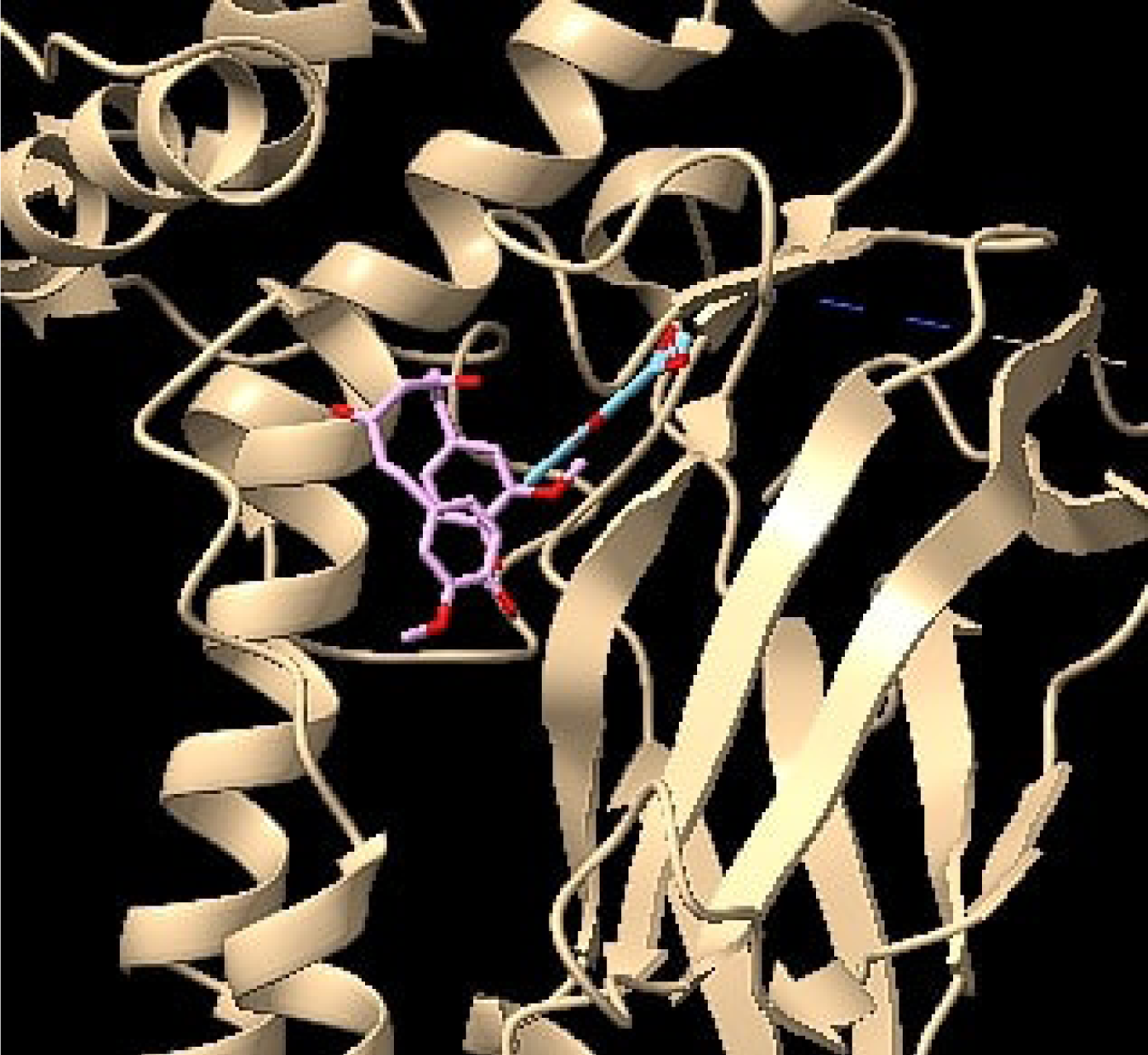
Close-up version of Figure 8 binding site. Lavender represents Piperine and Cyan represents Curcumin

**Fig. 10.**
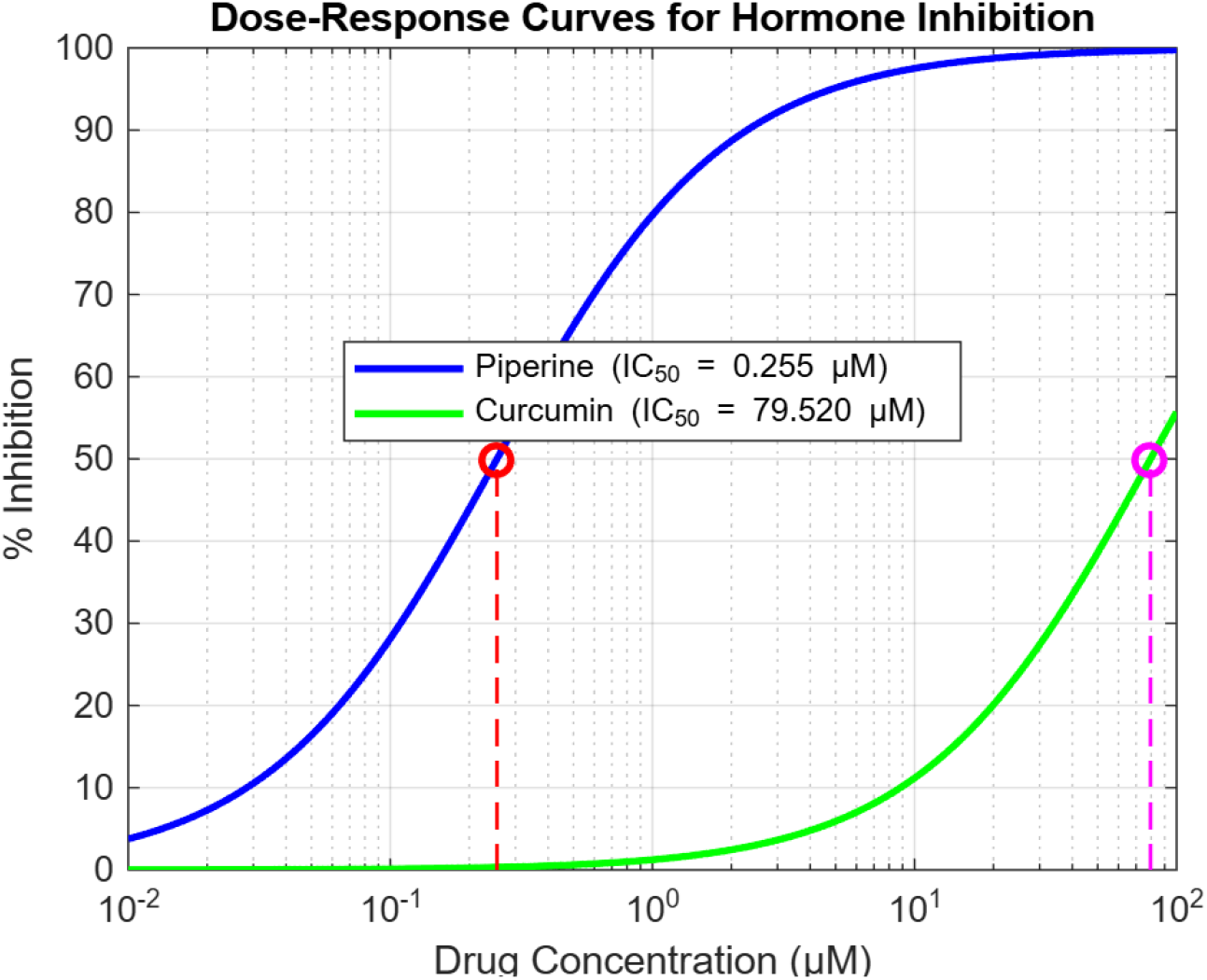
Monotherapy Curve of Curcumin (in green) and Piperine (in blue) with respect to the inhibition of TRPV1 protein.

**Fig. 11.**
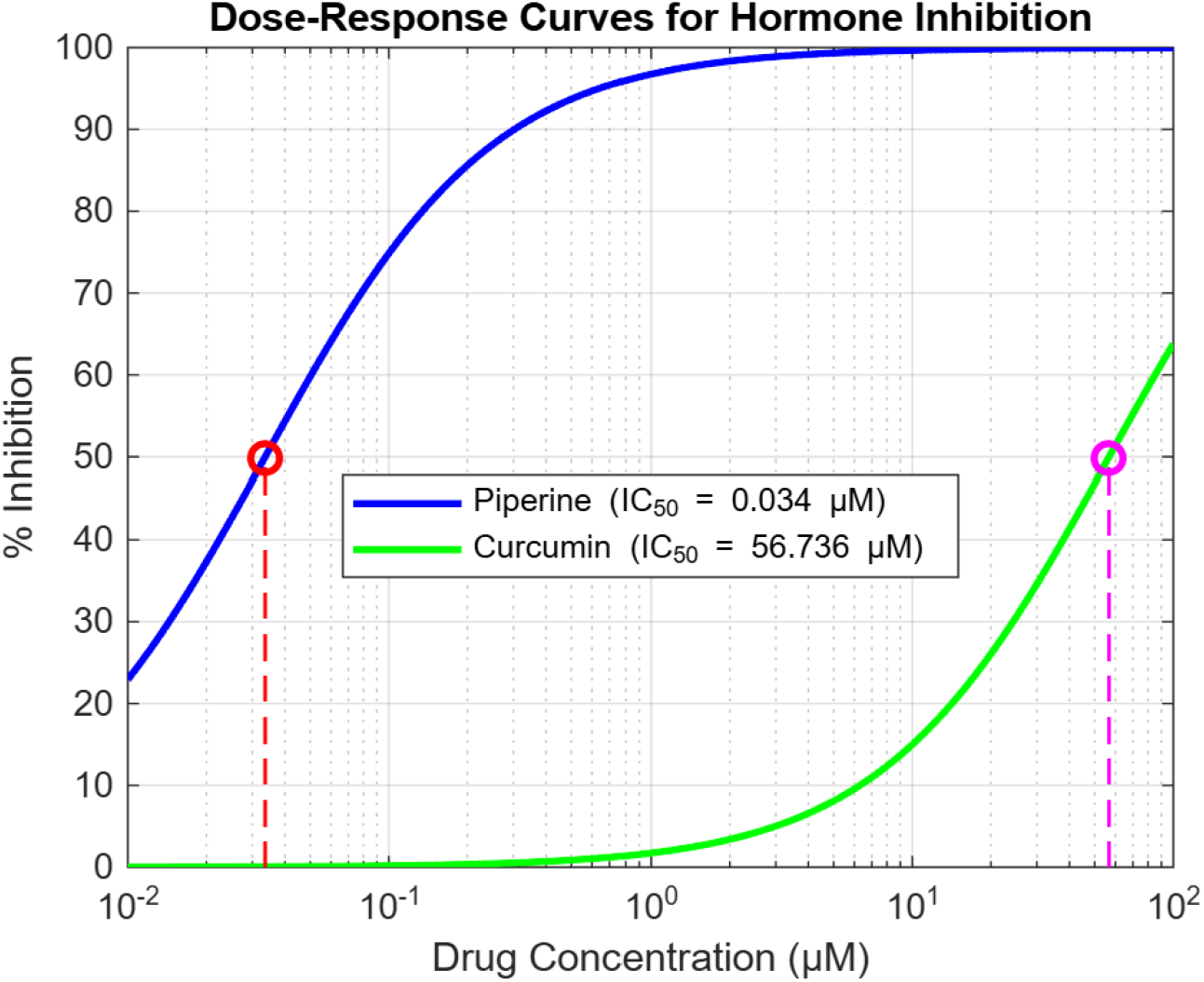
Monotherapy Curve of Curcumin (in green) and Piperine (in blue) with respect to the inhibition of STAT3 protein.

The large divergence of the *IC*_5_0 values infers that Piperine with a very low concentration can increase the combined effectiveness of Curcumin and itself. This is illustrated in figures 12 and 13. The plot is derived from the 4 parameter logistic curve of the Loewe’s Additivity Model mentioned in equation 5 and 6. In both the plots we range the shape parameter *λ* from 0 to 3 along the z - axis. This is done in order to fit with any potential experiments with a fixed shape parameter.

**Fig. 12.**
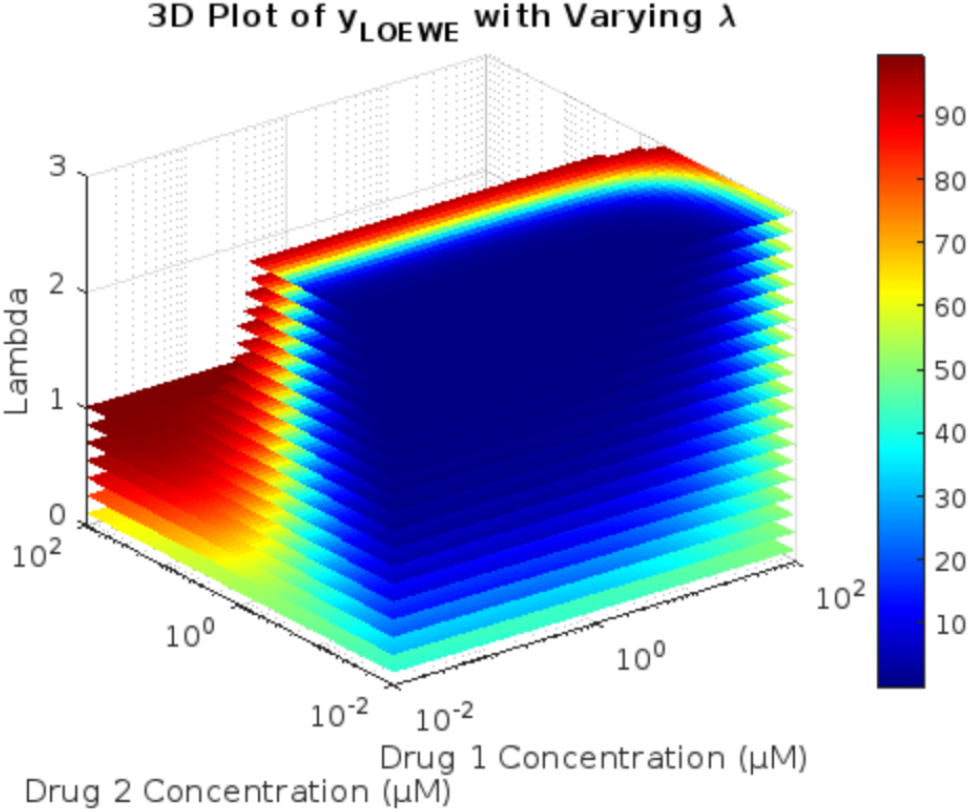
3D dose response matrix of combined effective inhibition of TRPV1 Protein by Curcumin and Piperine

**Fig. 13.**
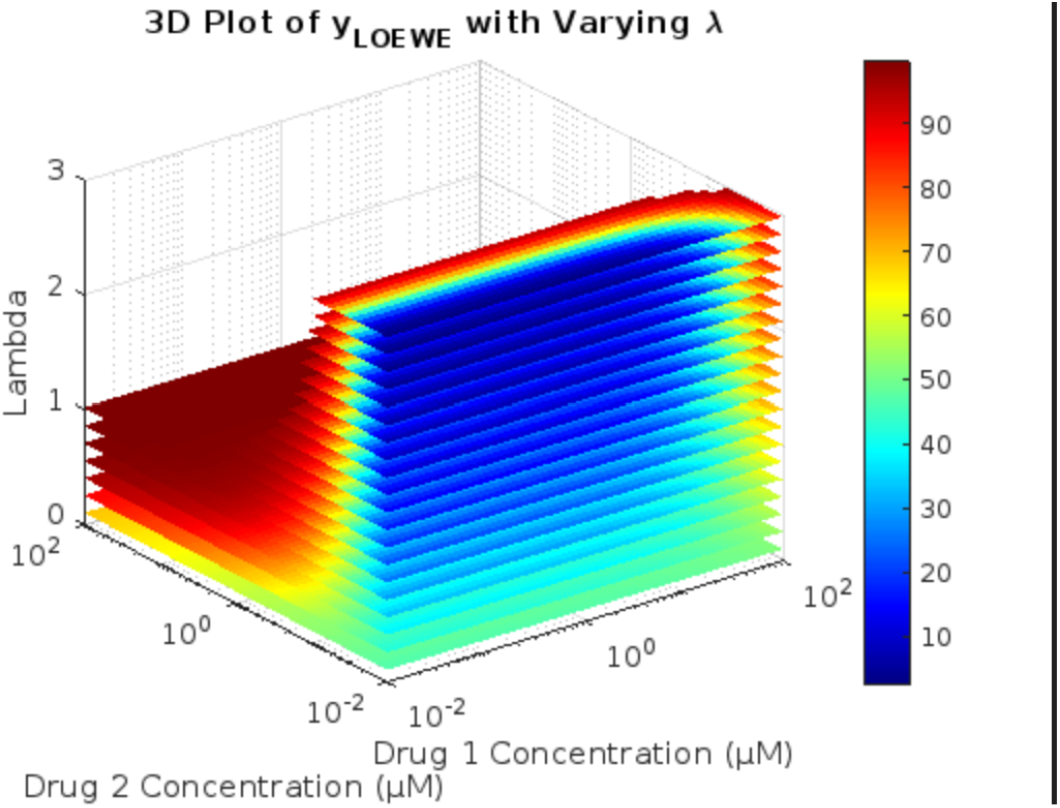
3D dose response matrix of combined effective inhibition of STAT3 Protein by Curcumin and Piperine

## 5 Conclusion

In this study, we have performed an in-silico analysis to validate and analyze the extent of synergetic properties of Curcumin and Piperine in the context of prevention and treatment of Breast Cancer. All computational experiments are done with the MCF7 cell kept in focus. We start with the prediction of the conventional Bliss drug synergy score using neural network regression and classification models. Both these models have predicted that the system of interest is synergetic in nature. We further analyze the extent of the synergy by Molecular Docking. We have used TRPV1 and STAT3 as our target proteins of interest. Using ChimeraX and PyRx, we have simulated simultaneous molecular docking of both ligands over the target protein and determined its potential binding affinity. Using FPocket software, we have further analyzed the properties of the binding site of the target protein where the ligands have docked. From the retrieved binding affinity values we have determined the dose response curves of both Curcumin and Piperine with respect to the inhibition percentage of the target proteins TRPV1 and STAT3. We have performed both independent monotherapy curves and 3D heatmap plot of inhibition due to the simultaneuous effect of both ligands on the basis of the Loewe’s Additivity Model. From this, we can infer that Piperine synergistically improves the effectiveness of Curcumin as substantiated from published literature. For our future work, we aim to simulate ADME Properties (Absorption, Distribution, Metabolism and Excretion) in order to prove bioavailability of the two components and also to work on virtual clinical trials and predict its effect in the human body in order to save time and energy on lab work and improve the safety of human clinical trials.

## Supplementary information

Code is available upon request.

## Acknowledgements

We would like to thank VIT Chennai and MedxAI Innovations Pvt. Ltd for their continued support towards the project.

## Declarations

Some journals require declarations to be submitted in a standardised format. Please check the Instructions for Authors of the journal to which you are submitting to see if you need to complete this section. If yes, your manuscript must contain the following sections under the heading ‘Declarations’:

- Funding
- The authors of this paper declare no conflict of interest.
- Ethics approval and consent to participate
- Consent for publication
- Data available upon request
- Materials availability
- Code available upon request.
- Author contribution

If any of the sections are not relevant to your manuscript, please include the heading and write ‘Not applicable’ for that section.

## Notes

### Competing Interest Statement

The authors have declared no competing interest.

## References

[1] Aggarwal, B.B., Sundaram, C., Malani, N., Ichikawa, H.: In: Aggarwal, B.B., Surh, Y.-J., Shishodia, S. (eds.) CURCUMIN: THE INDIAN SOLID GOLD, pp. 1–75. Springer, Boston, MA (2007)

[2] Gupta, S.C., Prasad, S., Kim, J.H., Patchva, S., Webb, L.J., Priyadarsini, I.K., Aggarwal, B.B.: Multitargeting by curcumin as revealed by molecular interaction studies. Natural product reports 28(12), 1937–1955 (2011)

[3] Maheshwari, R.K., Singh, A.K., Gaddipati, J., Srimal, R.C.: Multiple biological activities of curcumin: A short review. Life Sciences 78(18), 2081–2087 (2006) 10.1016/j.lfs.2005.12.007. NATURECEUTICALS (NATURAL PRODUCTS), NUTRACEUTICALS, HERBAL BOTANICALS, AND PSYCHOACTIVES: DRUG DISCOVERY AND DRUG-DRUG INTERACTIONS

[4] Huanbutta, K., Burapapadh, K., Kraisit, P., Sriamornsak, P., Ganokratanaa, T., Suwanpitak, K., Sangnim, T.: Artificial intelligence-driven pharmaceutical industry: A paradigm shift in drug discovery, formulation development, manufacturing, quality control, and post-market surveillance. European Journal of Pharmaceutical Sciences 203, 106938 (2024) 10.1016/j.ejps.2024.106938

[5] Jurenka, J.S.: Anti-inflammatory properties of curcumin, a major constituent of curcuma longa: a review of preclinical and clinical research. Alternative medicine review : a journal of clinical therapeutic 14 2, 141–53 (2009)

[6] Shoba, G., Joy, D., Joseph, T., Majeed, M., Rajendran, R.S., Srinivas, P.: Influence of piperine on the pharmacokinetics of curcumin in animals and human volunteers. Planta medica 64 4, 353–6 (1998)

[7] Goel, A., Kunnumakkara, A.B., Aggarwal, B.B.: Curcumin as “curecumin”: From kitchen to clinic. Biochemical Pharmacology 75(4), 787–809 (2008) 10.1016/j.bcp.2007.08.016

[8] Gupta, S.C., Kismali, G., Aggarwal, B.B.: Curcumin, a component of turmeric: From farm to pharmacy. BioFactors 39(1), 2–13 (2013) 10.1002/biof.1079 https://iubmb.onlinelibrary.wiley.com/doi/pdf/10.1002/biof.1079

[9] Li, H., Zou, L., Kowah, J.A., He, D., Wang, L., Yuan, M., Liu, X.: Predicting drug synergy and discovering new drug combinations based on a graph autoencoder and convolutional neural network. Interdisciplinary Sciences: Computational Life Sciences 15(2), 316–330 (2023)

[10] Schwarz, K., Pliego-Mendieta, A., Mollaysa, A., Planas-Paz, L., Pauli, C., Allam, A., Krauthammer, M.: Ddos: a graph neural network based drug synergy prediction algorithm. arXiv preprint arXiv:2210.00802 (2022)

[11] Malyutina, A., Majumder, M.M., Wang, W., Pessia, A., Heckman, C.A., Tang, J.: Drug combination sensitivity scoring facilitates the discovery of synergistic and efficacious drug combinations in cancer. PLOS Computational Biology 15(5), 1–19 (2019) 10.1371/journal.pcbi.1006752

[12] Luo, H., Yin, W., Wang, J., Zhang, G., Liang, W., Luo, J., Yan, C.: Drug-drug interactions prediction based on deep learning and knowledge graph: A review. Iscience (2024)

[13] Liu, X.-Y., Mei, X.-Y.: Prediction of drug sensitivity based on multi-omics data using deep learning and similarity network fusion approaches. Frontiers in Bioengineering and Biotechnology 11 (2023) 10.3389/fbioe.2023.1156372

[14] Landrum, G.: RDKit: Open-Source Cheminformatics Software. Accessed: 2023-11-13 (2023). 10.5281/zenodo.591637. https://www.rdkit.org

[15] Zeng, X., Xiang, H., Yu, L., Wang, J., Li, K., Nussinov, R., Cheng, F.: Accurate prediction of molecular targets using a self-supervised image representation learning framework. Research Square (2022)

[16] Zagidullin, B., Aldahdooh, J., Zheng, S., Wang, W., Wang, Y., Saad, J., Malyutina, A., Jafari, M., Tanoli, Z., Pessia, A., Tang, J.: Drugcomb: an integrative cancer drug combination data portal. Nucleic Acids Research 47(W1), 43–51 (2019) 10.1093/nar/gkz337 https://academic.oup.com/nar/article-pdf/47/W1/W43/28880110/gkz337.pdf

[17] Weininger, D.: Smiles, a chemical language and information system. 1. introduction to methodology and encoding rules. Journal of Chemical Information and Computer Sciences 28(1), 31–36 (1988) 10.1021/ci00057a005 https://doi.org/10.1021/ci00057a005

[18] Rogers, D., Hahn, M.: Extended-connectivity fingerprints. Journal of Chemical Information and Modeling 50(5), 742–754 (2010) 10.1021/ci100050t https://doi.org/10.1021/ci100050t. PMID: 20426451

[19] Agarap, A.F.: Deep learning using rectified linear units (relu). CoRR abs/1803.08375 (2018) 1803.08375

[20] Paszke, A., Gross, S., Massa, F., Lerer, A., Bradbury, J., Chanan, G., Killeen, T., Lin, Z., Gimelshein, N., Antiga, L., Desmaison, A., Köpf, A., Yang, E., DeVito, Z., Raison, M., Tejani, A., Chilamkurthy, S., Steiner, B., Fang, L., Bai, J., Chintala, S.: PyTorch: An Imperative Style, High-Performance Deep Learning Library (2019). https://arxiv.org/abs/1912.01703

[21] National Center for Biotechnology Information: PubChem Compound Summary for CID 969516, Curcumin. https://pubchem.ncbi.nlm.nih.gov/compound/Curcumin. Accessed 13 November, 2024

[22] National Center for Biotechnology Information: PubChem Compound Summary for CID 638024, Piperine. Retrieved November 13, 2024 from https://pubchem.ncbi.nlm.nih.gov/compound/Piperine (2024)

[23] O’Boyle, N.M., Banck, M.S., James, C.A., Morley, C., Vandermeersch, T., Hutchison, G.R.: Open babel: An open chemical toolbox. Journal of Cheminformatics 3, 33–33 (2011)

[24] Berman, H.M., Westbrook, J., Feng, Z., Gilliland, G., Bhat, T.N., Weissig, H., Shindyalov, I.N., Bourne, P.E.: The protein data bank. Nucleic Acids Research 28(1), 235–242 (2000) 10.1093/nar/28.1.235 https://academic.oup.com/nar/article-pdf/28/1/235/9895144/280235.pdf

[25] Meng, E.C., Goddard, T.D., Pettersen, E.F., Couch, G.S., Pearson, Z.J., Morris, J.H., Ferrin, T.E.: Ucsf chimerax: Tools for structure building and analysis. Protein Science 32(11), 4792 (2023) 10.1002/pro.4792 https://onlinelibrary.wiley.com/doi/pdf/10.1002/pro.4792

[26] Pettersen, E.F., Goddard, T.D., Huang, C.C., Meng, E.C., Couch, G.S., Croll, T.I., Morris, J.H., Ferrin, T.E.: Ucsf chimerax: Structure visualization for researchers, educators, and developers. Protein Science 30(1), 70–82 (2021) 10.1002/pro.3943 https://onlinelibrary.wiley.com/doi/pdf/10.1002/pro.3943

[27] Dallakyan, S., Olson, A.J.: Small-molecule library screening by docking with pyrx. Methods in molecular biology 1263, 243–50 (2015)

[28] Schmidtke, P., Le Guilloux, V., Maupetit, J., Tuffïry, P.: fpocket: online tools for protein ensemble pocket detection and tracking. Nucleic Acids Research 38, 582–589 (2010) 10.1093/nar/gkq383 https://academic.oup.com/nar/article-pdf/38/suppl 2/W582/3857546/gkq383.pdf

[29] Yung-Chi, C., Prsoff, W.H.: Relationship between the inhibition constant (ki) and the concentration of inhibitor which causes 50 per cent inhibition (i50) of an enzymatic reaction. Biochemical Pharmacology 22(23), 3099–3108 (1973)

[30] Ianevski, A., He, L., Aittokallio, T., Tang, J.: Synergyfinder: a web application for analyzing drug combination dose–response matrix data. Bioinformatics 33(15), 2413–2415 (2017) 10.1093/bioinformatics/btx162 https://academic.oup.com/bioinformatics/article-pdf/33/15/2413/50756595/bioinformatics 33 15 2413.pdf

[31] Yadav, B., Wennerberg, K., Aittokallio, T., Tang, J.: Searching for drug synergy in complex dose–response landscapes using an interaction potency model. Computational and Structural Biotechnology Journal 13, 504–513 (2015) 10.1016/j.csbj.2015.09.001

[32] Inc., T.M.: MATLAB Version: 9.13.0 (R2022b). https://www.mathworks.com

